# Context-aware geometric deep learning for RNA sequence design

**DOI:** 10.1101/2025.06.21.660801

**Authors:** Parth Bibekar, Lucien F. Krapp, Matteo Dal Peraro

## Abstract

RNA design has emerged to play a crucial role in synthetic biology and therapeutics. Although tertiary structure-based RNA design methods have been developed recently, they still overlook the broader molecular context, such as interactions with proteins, ligands, DNA, or ions, limiting the accuracy and functionality of designed sequences. To address this challenge, we present RISoTTo (RIbonucleic acid Sequence design from TerTiary structure), a parameter-free geometric deep learning approach that generates RNA sequences conditioned on both their backbone scaffolds and the surrounding molecular context. We evaluate the designed sequences based on their native sequence recovery rate and further validate them by predicting their secondary structures *in silico* and comparing them to the corresponding native structures. RISoTTo performs well on both metrics, demonstrating its ability to generate accurate and structurally consistent RNA sequences. Additionally, we present an *in silico* design study of domain 1 of the NAD^+^ riboswitch, where RISoTTo-generated sequences are predicted to exhibit enhanced binding affinity for both the U1A protein and the NAD^+^ ligand.

## 1 Introduction

RNA design plays a central role in synthetic biology, enabling the development of functional RNA molecules for a variety of applications, including gene expression control, biosensors, and nanotechnology [1, 2, 3]. RNA inverse folding refers to the design of RNA sequences that are predicted to fold into a target structural conformation. RNA design methods have traditionally focused on predicting secondary structures, a problem known to be NP-hard [4]. The foundational work by Hofacker et al. (1994) marked the beginning of systematic efforts to address this challenge, introducing an efficient local search strategy to optimize a seed sequence with an energy function [5]. Subsequently, numerous methods have been developed for secondary structure-based RNA design, relying on energy minimization guided by thermodynamic parameters [6, 7]. In parallel, probabilistic methods for modeling sequence distribution have been extensively investigated as alternative approaches [8, 9, 10]. More recently, various alternative optimization strategies have been proposed, including genetic algorithms [11], constraint programming [12] and ant colony optimization [13]. Building on these efforts, advances in deep learning have enabled the development of data-driven methods for RNA sequence design. For example, LEARNA and Meta-LEARNA Runge et al. [14] use reinforcement learning to generate sequences that fold into target secondary structures.

In contrast, three-dimensional RNA design has received comparatively less attention. Early methods like FARNA [15] and Rosetta [16] fixed-backbone redesign laid the groundwork for energy-based optimization in this space. More recently, advances in deep learning, along with the growing availability of high-quality RNA tertiary structure data, have enabled the development of data-driven methods for 3D RNA design. gRNAde employs a multi-state graph neural network combined with autoregressive decoding to design RNA sequences conditioned on one or more 3D backbone structures, allowing design across multiple conformations and capturing the intrinsic structural flexibility of RNA [17]. RiboDiffusion frames the RNA inverse folding problem as learning the sequence distribution conditioned on a fixed 3D backbone using a generative diffusion model [18]. RhoDesign is a structure-to-sequence deep learning platform designed for the de novo generative design of RNA aptamers [19]. It guides sequence generation through structural predictions, enabling the design of RNA sequences that are structurally similar to known aptamers. However, these methods primarily focus on designing RNA sequences to match a target 3D conformation in isolation, without accounting for the broader molecular context in which RNAs often function. RNA structures are frequently stabilized or altered through interactions with proteins [20], small molecules [21], ions [22], or even DNA [23], making it essential to account for these interactions when designing functional RNA sequences intended for use in biological contexts.

Techniques from protein modeling are frequently adapted to RNA, given that protein design has long been a central focus in computational biology. A recent advancement in this space is CARBonAra (Context-aware Amino acid Recovery from Backbone Atoms and heteroatoms), a protein design model designed to recover sequences that fit a given backbone scaffold [24]. Built upon the Protein Structure Transformer (PeSTo) framework [25], CARBonAra utilizes a geometric transformer architecture that operates directly on atomic coordinates and element types, without requiring any structural preprocessing. A key strength of the model is its context-awareness; it integrates molecular interactions with ligands, nucleic acids, ions, and other proteins into its predictions. This makes CARBonAra particularly effective at recovering functional sequences in biologically relevant environments. Given its flexible architecture and ability to incorporate molecular context, it makes CARBonAra a natural candidate for adaptation to context-aware RNA sequence design.

We adapt the CARBonAra framework to introduce RISoTTo (RIbonucleic acid Sequence design from TerTiary structure), a parameter-free geometric transformer model for RNA sequence design conditioned on fixed tertiary structures. RISoTTo explicitly incorporates interactions with non-RNA entities including proteins, small molecules, ions, and DNA, enabling context-aware design. The model is trained and validated on RNA-containing tertiary structures from the PDB database [26], as well as on predicted structures generated using trRosettaRNA [27]. We evaluate RISoTTo on a benchmark test set to assess native sequence recovery and validate the designability of its outputs through *in silico* secondary and tertiary structure prediction using EternaFold [28] and AlphaFold 3 [29], respectively. Additionally, we present an *in silico* design study of domain 1 of the NAD^+^ riboswitch, demonstrating RISoTTo’s capacity to generate sequences predicted to exhibit favorable interactions with both its cognate protein partner U1A and its ligand NAD^+^ [30]. We hope that RISoTTo will serve as a useful tool for advancing RNA design, enabling the design of functionally relevant RNA sequences that consider both structural constraints and molecular context, ultimately contributing to the development of RNA-based therapeutics.

## 2 Methods

At the core of RISoTTo is a geometric transformer model, operating on molecular complexes represented as atomic point clouds, consisting of both the RNA backbone scaffold and surrounding non-RNA atoms. Each atom in the complex is labeled with its atomic element type and is associated with a scalar state (*q*) and a vector state (***p***), while the geometry is encoded through pairwise distances (*D*) and normalized displacement vectors (***R***). Each layer (*l*) of the geometric transformer (***GT***) processes these features using progressively larger neighborhoods, starting with the 8 nearest neighbors (*nn*) and increasing up to 64, as described in **eq. 1**.

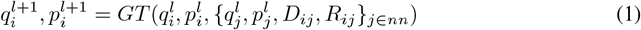

Here, *D*_*ij*_ ∈ ℝ denotes the Euclidean distance between atoms *i* and *j*, and **R**_*ij*_ ∈ ℝ^3^ is the normalized displacement vector from *i* to *j*. The geometric transformer in RISoTTo implements an attention mechanism based on the standard query–key-value approach [31] as formalized in **eq. 2** and **3**. For each atom *i*, the scalar and vector queries *Q*_*q*_ and *Q*_*p*_ are computed as functions of its current scalar and vector states 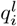 and 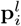, respectively. The keys (*K*) are derived from the set of neighboring atoms *j ∈ nn*_*i*_, and is a function of the local interaction features 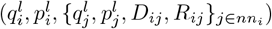.

Subsequently, scalar value vectors (*V*_*q*_) and vector value vectors (*V*_*p*_) are extracted from the computed scalar and vector quantities of these interactions. The transformer performs a linear combination of the vector features and states, ensuring that the resulting vector state is equivariant to rotation. Attention is computed over multiple heads, with the scalar and vector tracks projected using learned weights 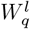 and 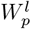, respectively.

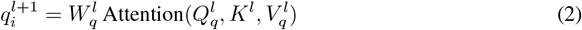

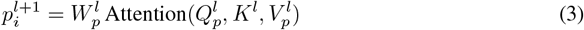

### Deep learning architecture

The architecture of RISoTTo closely follows that of CARBonAra [24]. One-hot encodings of the atomic element types are projected into a 64-dimensional scalar state using a three-layer neural network with hidden layers of size 64. This representation is then processed by a sequence of 20 geometric transformer layers, each with a key size of 3 and two attention heads, operating over neighborhoods ranging from 8 to 64 nearest neighbors and residual connections are employed within each transformer layer, as illustrated in Figure 1. The atomic-level features within each nucleotide are then aggregated into a nucleotide-level representation using geometric residue pooling with a four-headed self-attention mechanism. This residue-level encoding is then passed through a sequence module comprising a multilayer perceptron (MLP) with a hidden size of 64, which outputs a confidence score for each nucleotide via a sigmoid activation ranging from 0 to 1. These confidence scores can be interpreted as a probability estimate by modeling the likelihood of a correct prediction given a confidence score for each nucleotide type (see Supplementary Figure 1). This allows us to construct a final position weight matrix (PWM) from which RNA sequences can be sampled.

**Figure 1:**
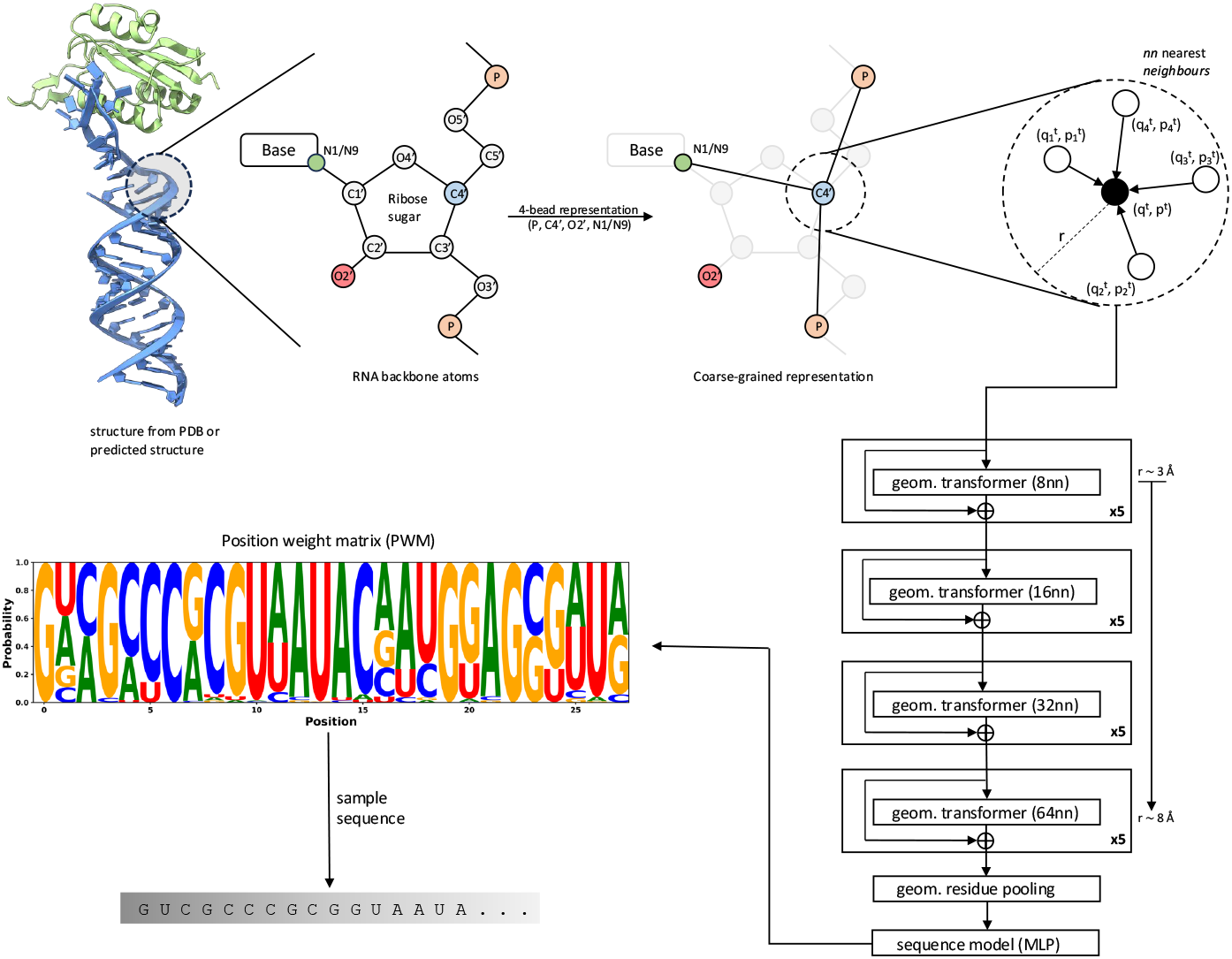
Overview of RISoTTo workflow for context-aware RNA design. The RNA backbone is described using a 4-bead coarse-grained representation, capturing the coordinates of four atoms per nucleotide: P, C4’, O2’, and either N1 (for pyrimidines) or N9 (for purines). The model comprises 20 layers of geometric transformers with residual connections, progressively expanding the neighborhood size from 8 to 64 nearest neighbors. Structural information is aggregated into a residue-level representation through transformer-based geometric pooling. The residue-level representations are aggregated and finally passed through a multilayer perceptron (MLP) to generate the final position weight matrix (PWM).

### Dataset

To train RISoTTo, we compiled RNA-containing subunits from the Protein Data Bank (PDB) [26], including standalone RNAs, protein–RNA complexes, and RNA–DNA hybrid complexes. For larger RNA subunits whose structures are not available in the legacy PDB format, we retrieved the corresponding entries from the RNASolo database [32]. To enable fair comparison with existing methods, we adopted the dataset split defined by Joshi et al. [17], which was used to train and evaluate gRNAde. This results in a training set of approximately 14,000 RNA subunits, a validation set of 693 subunits, and a test set of 379 subunits. The split ensures structural dissimilarity between sets using a TM-score threshold of <0.45, thereby preventing data leakage. The test set also includes 14 high-quality RNAs curated by Das et al. [16], which we use as an additional benchmark for RISoTTo. The distribution of sequence lengths across the dataset is shown in Supplementary Figure 3.

To further improve model generalization, we augmented the training set with approximately 5.5k RNA subunits by predicting tertiary structures for sequences from the bpRNA database [33] using trRosettaRNA [27]. These sequences, which have experimentally determined secondary structures, were originally used for self-distillation in training trRosettaRNA. To ensure that this augmentation enhances structural diversity rather than introducing redundancy, we applied two filtering criteria: (1) less than 80% sequence identity to any sequence in the full dataset (train, validation, and test) using CD-HIT [34], and (2) a TM-score below 0.45 with respect to the test and validation sets using US-align [35]. This ensures that the additional structures introduce novel folds and sequence contexts. After augmentation, the final training set consists of approximately 19.5k RNA subunits.

### Features and labels

We represent each nucleotide in the RNA backbone using a 4-bead coarse-grained model comprising the atoms P, C4’, O2’, and either N1 (for pyrimidines) or N9 (for purines). The subset of atoms P, C4’, and N1/N9 corresponds to the well-established “pseudotorsional” representation, which is sufficient to describe RNA backbone conformations in most cases [36]. However, since RISoTTo performs context-aware RNA design by incorporating information from neighboring atoms, we further analyzed local atomic contacts within a 5 Å radius around each backbone atom across the training set. This analysis revealed that the O2’ atom exhibits the highest frequency of contacts, suggesting its importance in local structural environments (see Figure 1 and Supplementary Figure 2). Moreover, the O2’ atom is located in the shallow groove of RNA, a known hotspot for RNA–protein interactions and hydration sites in many RNAs [37]. Based on both empirical and structural insights, we included O2’ in our backbone representation to better capture the local context critical for functional design.

The input scalar features are one-hot encodings of the 30 most common atomic elements, while geometric context is captured through pairwise distances and normalized displacement vectors between atoms; the associated vector states are initialized by sampling from an isotropic normal distribution. RISoTTo requires no additional physicochemical descriptors or handcrafted features and can operate even on incomplete molecular structures.

### Training

We trained the model for 48 hours on NVIDIA H100 GPUs (64 GB memory). To manage memory constraints, we excluded structures with more than 12,288 atoms and filtered out RNAs shorter than 20 nucleotides. The model was optimized using the Adam optimizer [38] with a learning rate of 1 × 10^−4^ and binary cross-entropy (BCE) as the loss function [39]. To address nucleotide imbalance in the dataset, we applied class weighting based on the nucleotide distribution.

### Evaluation

We evaluate our designed sequences on the test set curated by Joshi et al. [17], which also includes 14 high-quality structured RNAs selected by Das et al. [16]. On this additional benchmark, we compare RISoTTo’s native sequence recovery against several baseline methods: ViennaRNA (2D design only) [40], FARNA [15], Rosetta [16], RDesign [41], and gRNAde [17]. For the full test set, we report native sequence recovery accuracy.

To assess structural consistency, we compute 2D structure consistency by folding the designed sequences using EternaFold [28] and calculating the average Matthews Correlation Coefficient (MCC) with respect to the native secondary structures. For the 14 benchmark RNAs from Das et al., we also evaluate 3D structural consistency by predicting their 3D structures using AlphaFold 3 [29] and computing RMSD, GDT_TS, and TM-score against the native structures.

## 3 Results and Discussions

We set out to compare RISoTTo’s performance against existing RNA design methods by evaluating sequence recovery with several baseline models, including ViennaRNA (2D design only) [40], FARNA [15], Rosetta [16], RDesign[41], and gRNAde[17]. On a benchmark set of 14 structured RNAs curated by Das et al. [16], RISoTTo achieves the highest average sequence recovery of 61%, outperforming gRNAde (56.8%), RDesign (43.0%), Rosetta (45.0%), FARNA (32.1%), and ViennaRNA (26.9%), as shown in Figure 2a. On the larger test set from Joshi et al. [17] of 379 subunits, RISoTTo achieves an average sequence recovery of 59.3%, compared to 54.3% for gRNAde, indicating a modest yet consistent improvement (Figure 2b). We also assess 2D structural consistency by computing the self-consistency Matthews Correlation Coefficient (sc2D MCC) between the EternaFold [28] predicted and native secondary structures, which yields an average score of 0.43 (Figure 3a).

**Figure 2:**
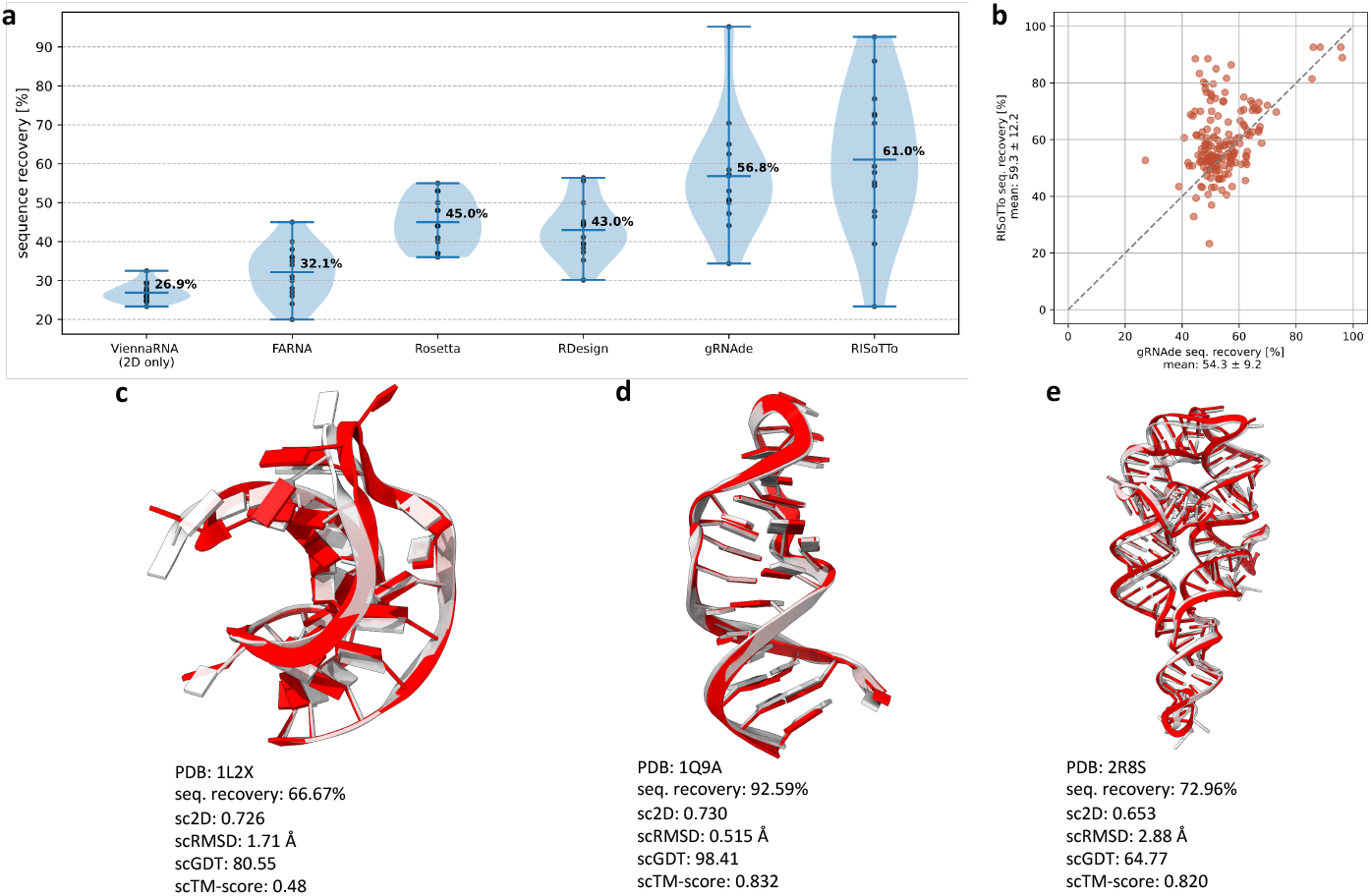
Benchmarking RISoTTo against existing RNA design methods. (a) Sequence recovery on a benchmark set of 14 structured RNAs [16], comparing RISoTTo to baseline methods including ViennaRNA [40], FARNA [15], Rosetta [16], RDesign [41], and gRNAde [17] RISoTTo achieves the highest average recovery (62.0%). (b) Native sequence recovery of RISoTTo vs gRNAde on a larger test set from Joshi et al. [17] (c–e) Cherry-picked examples where AlphaFold 3 predicted tertiary structures from RISoTTo-designed sequences closely match the native folds.

**Figure 3:**
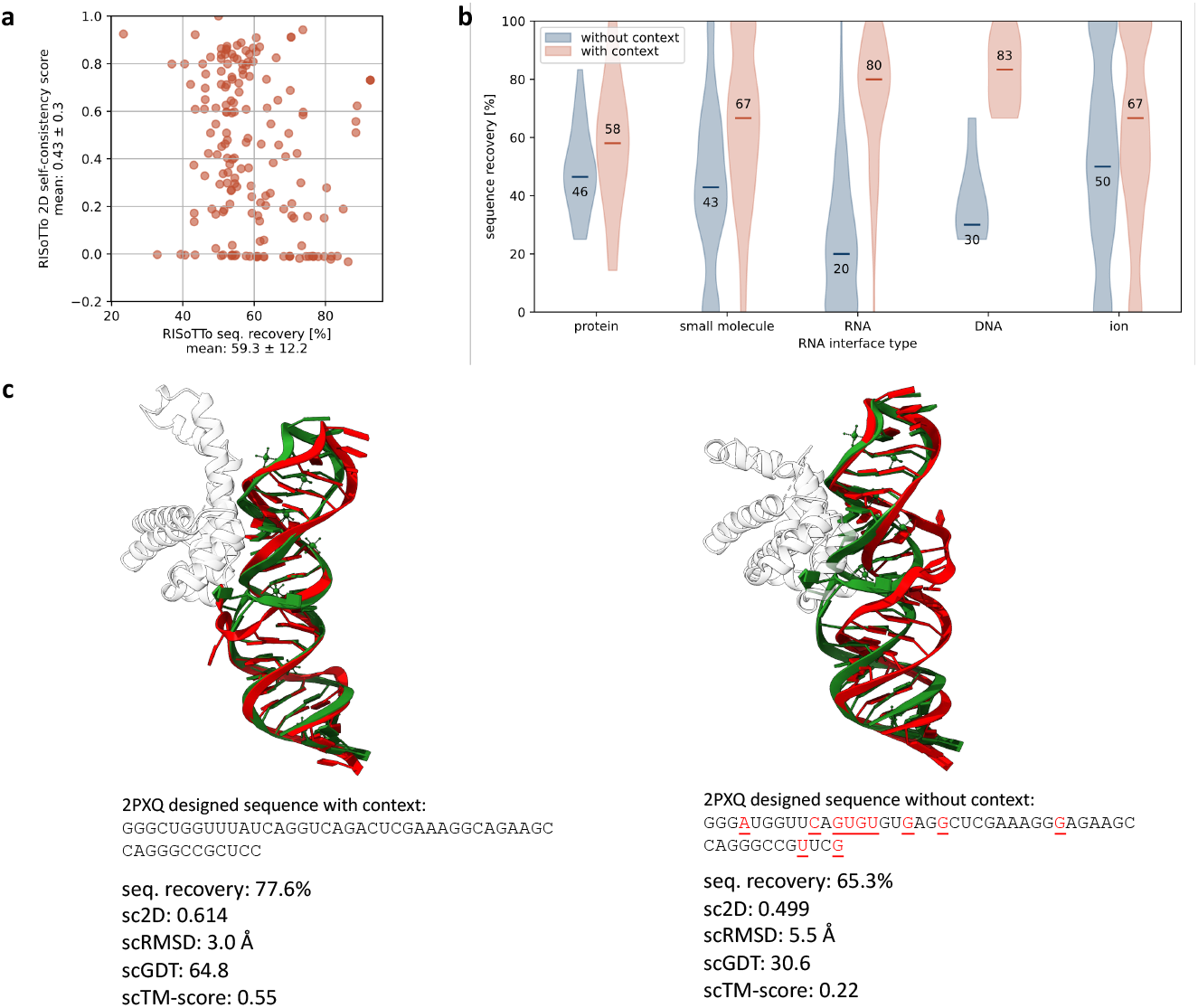
Evaluation of RISoTTo’s context-aware RNA design performance. (a) Native sequence recovery versus secondary structure consistency (sc2D) across the test set. (b) Comparison of sequence recovery at nucleotide interfaces with and without molecular context for different interaction types, including protein, small molecule, RNA, DNA, and ion. (c) Predicted tertiary structures for the *E. coli* SRP RNA complex designed with (left) and without (right) its ribonucleoprotein. Red indicates the predicted structure and green the native structure. Change in sequence prediction without context is highlighted.

We further highlight our designs through representative examples by aligning the tertiary structures predicted using AlphaFold 3 from RISoTTo-designed sequences with their corresponding native structures. These include the viral RNA pseudoknot from beet western yellow virus (PDB 1L2X, Figure 2c) [42], the sarcin/ricin domain from E. coli 23S rRNA (PDB 1Q9A, Figure 2d) [43], and a synthetic FAB bound to the P4–P6 RNA ribozyme domain (PDB 2R8S, Figure 2e) [44]. We take the TM-score, RMSD, and GDT score of the C4’ backbone atoms to measure the similarity between the predicted RNA structures of generated sequences and the given native backbones, demonstrating RISoTTo’s ability to generate sequences that maintain the native fold.

We assess sequence recovery at nucleotides within 5 Å of interacting ligands to evaluate RISoTTo’s ability to design RNA sequences in diverse molecular environments. We compare performance on the same RNA backbones with and without incorporating ligand interactions into the input. Sequence recovery improves with ligand context across all interface types: for RNA–protein complexes from 46% (without protein context) to 58% (with protein context), 43% to 67% for RNA–small molecule, 20% to 80% for RNA–RNA, 30% to 83% for RNA–DNA, and 50% to 67% for RNA–ion interactions. These improvements suggest that RISoTTo captures the geometric and chemical constraints imposed by interacting partners, enabling more accurate and context-aware RNA sequence design (Figure 3b).

As a concrete example of RISoTTo’s ability to incorporate molecular context, we design variant 14 of the *E. coli*. SRP RNA in complex with its ribonucleoprotein core (PDB ID: 2PXQ) [45]. We perform two separate designs: one with the protein context included and one without. As shown in Figure 3c, the design with protein context achieves a sequence recovery of 77.6%, compared to 65.3% without context. Interestingly, despite both designs having relatively similar sequence recoveries, their structural consistencies differ significantly. When we predict secondary structures using EternaFold [28], the design with context yields a 2D structural consistency score (sc2D) of 0.61, indicating strong agreement with the native secondary structure. In contrast, the design without context scores of 0.49, suggesting a slightly more dissimilar fold. We observe a similar trend in tertiary structure prediction using AlphaFold 3 [29]. The structure predicted from the context-aware sequence shows an RMSD of 3.0 Å relative to the native structure, whereas the structure from the context-free sequence has a higher RMSD of 5.5 Å. Correspondingly, the structural similarity metrics improve from 30.6 to 64.8 in scGDT and from 0.22 to 0.55 in scTM-score when the molecular context is included. These results further demonstrate that RISoTTo benefits from the inclusion of interacting molecular context, particularly in capturing native-like folds.

Similar to CARBonAra [24] and gRNAde [17], RISoTTo also supports autoregressive sequence generation, enabling the creation of diverse sequence variants from a given RNA tertiary structure. To achieve this, RISoTTo can imprint prior sequence information onto the RNA backbone. This allows the model to condition future predictions on previously predicted or known nucleotides.

As an example, we generated 20 sequences for Domain 1 of the NAD^+^ riboswitch (PDB: 7D7V) [30]. First, we used RISoTTo to predict a position weight matrix (PWM) from the backbone structure. We then sampled an initial sequence by selecting the nucleotide with the highest confidence at each position. For the next round, we randomly imprinted 50% of this sequence back into the model, fixing these positions during prediction, while resampling the remaining positions. Repeating this process enables RISoTTo to explore diverse sequences while anchoring to high-confidence predictions, maintaining overall sequence quality.

Domain 1 of the NAD^+^ riboswitch interacts with both a ligand (NAD^+^) and a protein partner (U1A). We folded all 20 designed sequences using AlphaFold 3 with both binding partners present. To evaluate binding strength, we used RSAPred [46] to predict RNA–NAD^+^ affinity and PRA-Pred [47] to assess RNA–U1A affinity. Out of the 20 sequences, 10 showed improved binding to NAD^+^, and 2 showed improved binding to U1A compared to the wild type (WT). Notably, two sequences outperformed the WT on both fronts, with predicted binding affinities of –9.70 and –9.55 kcal/mol to U1A (WT: –9.50 kcal/mol), and –6.04 and –8.83 kcal/mol to NAD^+^ (WT: –5.31 kcal/mol). Figure 4a shows one of the designed sequences folded with AlphaFold 3, aligned with the native structure. The prediction conserves both the overall fold and key binding interactions of the native RNA. Figure 4b presents the distribution of predicted binding affinities for the 20 designed sequences. As an additional validation, we used RibonanzaNet [48] to predict chemical mapping reactivities for both the designed and native RNA sequences. RibonanzaNet was trained on experimental datasets where RNA sequences were synthesized and probed using chemical mapping techniques, specifically dimethyl sulfate (DMS) and SHAPE reagents such as 2A3. We predicted per-nucleotide reactivity profiles for both the native and designed sequences under 2A3 and DMS conditions. As shown in Figure 4c, the designed sequence closely mirrored the native reactivity landscape, achieving mean absolute errors (MAE) of 0.19 for 2A3 and 0.15 for DMS. This example highlights how RISoTTo’s imprinting-based sampling strategy can generate diverse RNA sequences and can be adapted flexibly for a variety of design scenarios.

**Figure 4:**
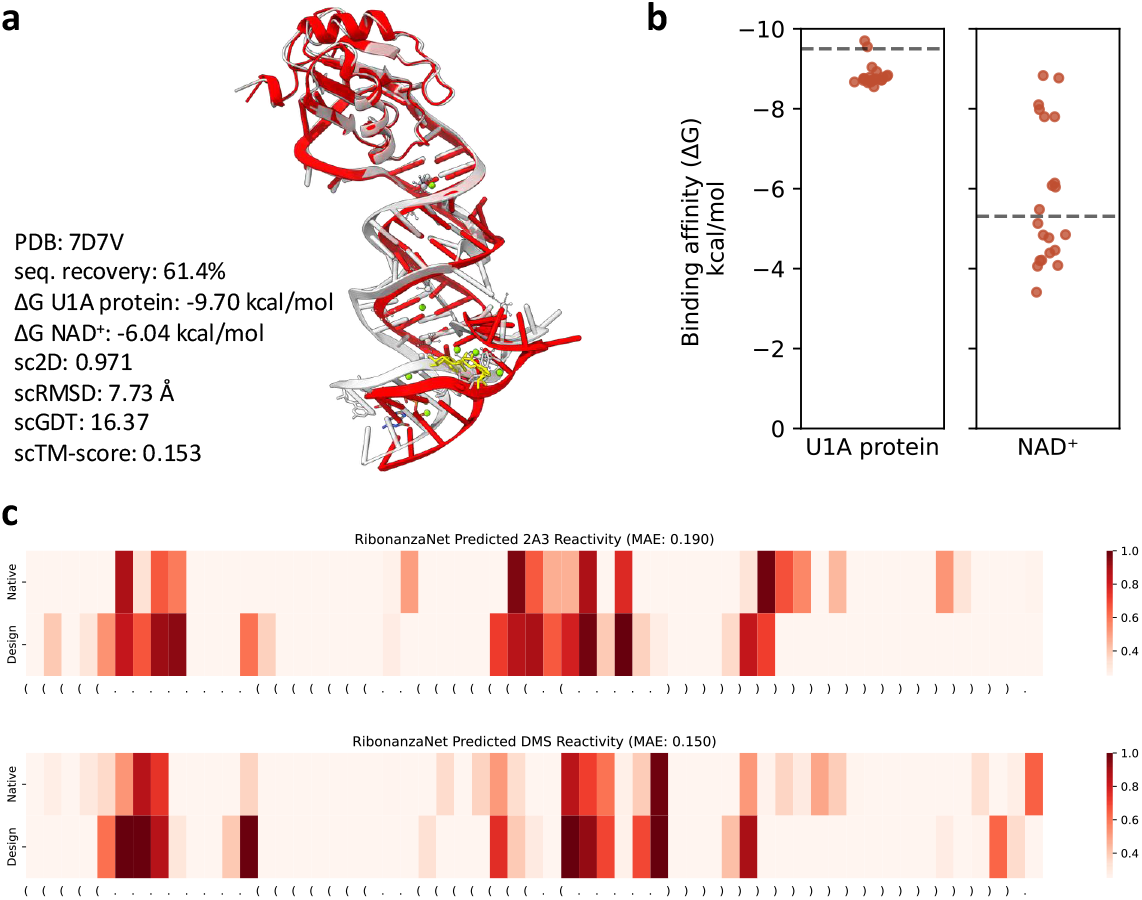
In silico design of the NAD^+^ riboswitch using RISoTTo. (a) Structural overlap of the RISoTTo-designed sequence with the native structure of Domain 1 of the NAD^+^ riboswitch (PDB: 7D7V), predicted using AlphaFold 3. (b) Distribution of predicted binding affinities for 20 autoregressively generated sequences with NAD^+^ (via RSAPred) and U1A protein (via PRA-Pred), dashed lines indicate predicted wild type binding affinities. (c) Predicted chemical mapping reactivity profiles for native and RISoTTo-designed RNA sequences using RibonanzaNet under 2A3 and DMS conditions.

## 4 Conclusion

We introduced RISoTTo, a context-aware model for RNA sequence design that leverages tertiary structure information. Benchmarking against recent methods like gRNAde shows a consistent improvement in native sequence recovery, demonstrating the advantage of incorporating molecular context, a limitation that has existed in previous approaches. We also demonstrate a practical use case where RISoTTo-generated sequences exhibit improved binding affinities, showcasing the model’s potential for guided design. In the future, we aim to explore more complex design scenarios, further expanding the applicability of RISoTTo to diverse RNA functionalities. While the RNA design field has seen rapid progress, the overall performance still lags behind protein inverse folding methods, likely due to the limited availability of high-quality RNA 3D structural data for training. Continued improvements in structure prediction methods and access to higher-quality RNA structure data will be crucial for advancing the field.

## Availability

All datasets, methods, and the RISoTTo source code presented in this work is available on github at https://github.com/LBM-EPFL/RISoTTo.

## Acknowledgments

The Swiss National Science Foundation is acknowledged for supporting this work (grant number 205321_192371 and 320030_232049 to MDP). We also acknowledge the Swiss National Supercomputing Centre (CSCS) for the generous computing time.

## Author contributions

PB and MDP conceived and designed the research project. PB implemented the RISoTTo code. PB, LFK and MDP analysed the results and wrote the paper.

## Supplementary Information

**Supplementary Table 1:** Performance comparison on 14 RNA structures curated by Das et al. [16]. Sequence recovery values for ViennaRNA, FARNA, Rosetta, and RDesign are taken from Joshi et al. [17]. gRNAde and RISoTTo report both sequence recovery and secondary structure self-consistency (sc2D) scores.

| PDB ID | Vienna | FARNa | Rosetta | RDesign | gRNade |  | RISoTTo |  |
| --- | --- | --- | --- | --- | --- | --- | --- | --- |
|  | Recovery | Recovery | Recovery | Recovery | Recovery | sc2D | Recovery | sc2D |
| 1CSL | 0.257 | 0.200 | 0.440 | 0.446 | **0.472** | 0.569 | 0.464 | **0.756** |
| 1ET4 | 0.259 | 0.340 | 0.440 | 0.393 | **0.625** | **-0.014** | 0.394 | -0.002 |
| 1F27 | 0.295 | 0.360 | 0.370 | 0.301 | **0.441** | 0.043 | 0.233 | **0.924** |
| 1L2X | 0.245 | 0.450 | 0.480 | 0.373 | 0.584 | 0.138 | **0.703** | **0.911** |
| 1LNT | 0.325 | 0.270 | 0.530 | 0.556 | 0.572 | **0.864** | **0.727** | -0.018 |
| 1Q9A | 0.275 | 0.400 | 0.410 | 0.442 | **0.952** | **0.777** | 0.926 | 0.731 |
| 4FE5 | 0.292 | 0.280 | 0.360 | 0.411 | **0.500** | **0.663** | 0.477 | 0.508 |
| 1X9C | 0.262 | 0.310 | 0.500 | 0.397 | 0.650 | -0.025 | **0.767** | **-0.011** |
| 1XPE | 0.271 | 0.240 | 0.400 | 0.383 | 0.530 | **0.912** | **0.543** | 0.825 |
| 2GCS | 0.249 | 0.260 | 0.440 | 0.452 | 0.508 | 0.406 | **0.577** | **0.470** |
| 2GDI | 0.250 | 0.380 | 0.480 | 0.352 | **0.569** | **0.728** | 0.551 | 0.283 |
| 2OEU | 0.233 | 0.300 | 0.370 | 0.500 | 0.504 | 0.060 | **0.593** | **0.241** |
| 2R8S | 0.269 | 0.360 | 0.530 | 0.564 | 0.704 | **0.778** | **0.723** | -0.003 |
| 354D | 0.280 | 0.350 | 0.550 | 0.446 | 0.344 | **0.452** | **0.863** | -0.033 |

**Supplementary Figure 1:**
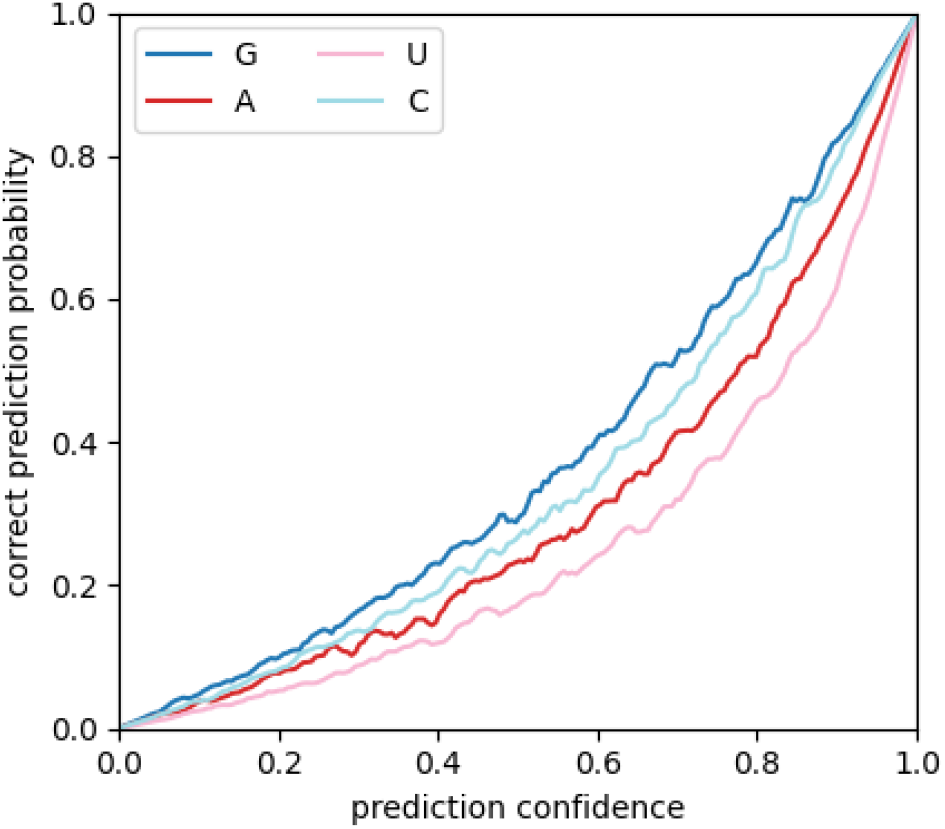
Relationship between prediction confidence and the prediction accuracy for each nucleotide type over the train set.

**Supplementary Figure 2:**
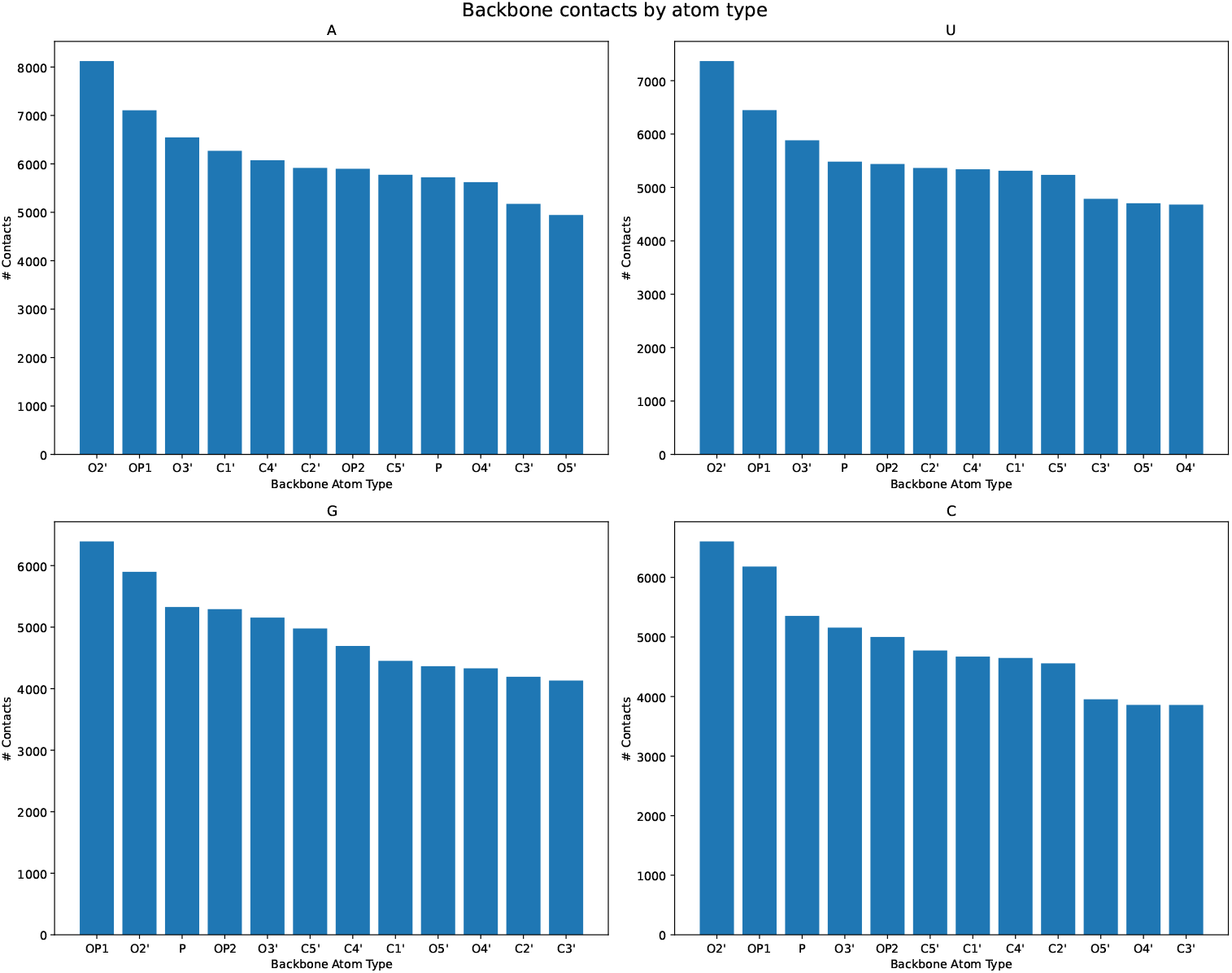
Distribution of atomic contacts within 5 Å for each backbone atom across all nucleotides. The O2’ atom exhibits a notably higher number of contacts.

**Supplementary Figure 3:**
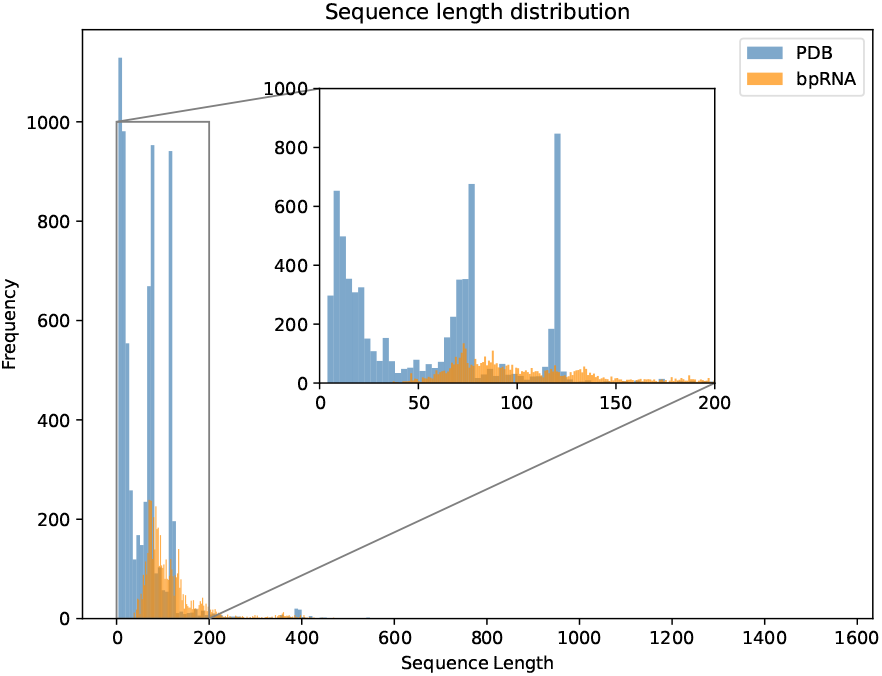
Distribution of RNA sequence lengths from the dataset (PDB in blue, bpRNA in orange).

